# Glutamylation imbalance leads to photoreceptor cell degeneration

**DOI:** 10.1101/2024.04.09.588666

**Authors:** Olivier Mercey, Sudarshan Gadadhar, Maria M. Magiera, Laura Lebrun, Corinne Kostic, Alexandre Moulin, Yvan Arsenijevic, Carsten Janke, Paul Guichard, Virginie Hamel

## Abstract

The stereotypic structure of microtubules, assembled from conserved α/β-tubulin dimers is subject to a complex diversity of Post-translational Modifications (PTMs). PTMs are predicted to fine-tune microtubule properties and interactions with other proteins, thus allowing microtubules to perform specific functions. Cilia accumulate several types of tubulin PTMs, such as polyglutamylation, polyglycylation, detyrosination and acetylation, whose functions are not yet fully understood. Recently, mutations of *AGBL5*, coding for the deglutamylating enzyme CCP5, have been associated to retinitis pigmentosa, suggesting that perturbation of polyglutamylation leads to the degeneration of photoreceptor cells. However, the molecular mechanisms underlying this degeneration remain unknown. Here, using super-resolution Ultrastructure Expansion Microscopy in mouse and human photoreceptor cells, we found that most tubulin PTMs are accumulated at the level of the connecting cilium, a structure linking the outer and inner segments of photoreceptor cells. Using mouse models with increased glutamylation (*Ccp5*^-/-^ and *Ccp1*^-/-^), or loss of tubulin acetylation (*Atat1*^-/-^), we demonstrated that aberrant glutamylation, but not loss of acetylation, resulted in perturbed molecular architecture of the outer segment, with the loss of the bulge region and destabilization of the distal axoneme. Concurrently, we observed a substantial impairment in tubulin glycylation and intraflagellar transport. Altogether our results indicate that glutamylation plays a crucial role in the maintenance of the molecular architecture of the outer segment and point to tubulin PTM imbalance as possible culprit in retinal degeneration.

## Introduction

Cilia are highly conserved organelles present on the surface of most eukaryotic cells. They exhibit remarkable structural complexity organized around an axoneme composed of nine microtubule doublets. Tubulin, the constituent of these microtubules, undergoes a diverse array of PTMs including acetylation, detyrosination, polyglutamylation, polyglycylation, phosphorylation, polyamination, SUMOylation, glycosylation, arginylation, methylation or palmitoylation. These modifications, referred to as one aspect of the tubulin code, are thought to influence on microtubule dynamics, stability, and interactions with associated proteins (Janke & Magiera, 2020). PTMs being mostly enriched on stable microtubules, cilia represent an interesting model to study these modifications. Furthermore, tubulin PTMs have emerged as key regulators of ciliary assembly, maintenance, and signaling (Yang et al., 2021).

Polyglutamylation, one of the most abundant PTMs in cilia (Yang et al., 2021) is probably the most studied tubulin PTM in these organelles. In motile cilia, it has been shown that glutamylation controls the activity of inner arm dynein, important for the regulation of ciliary beating (Kubo, Yanagisawa, Yagi, Hirono, & Kamiya, 2010; Suryavanshi et al., 2010). Recently, Alvarez Viar and colleagues revealed that polyglutamylation of protofilament 9 of the B-tubule is a conserved feature of motile cilia shared between algae and mice (Viar, Klena, Martino, Nievergelt, & Pigino, 2023). This highly localized distribution of this PTM allows for the interaction with the nexin-dynein regulatory complex (NDRC), and thus regulating ciliary beating behavior. Interestingly, both hypoglutamylation (Grau et al., 2013; Ikegami, Sato, Nakamura, Ostrowski, & Setou, 2010; Kubo et al., 2010; Pathak, Austin, & Drummond, 2011; Suryavanshi et al., 2010) and hyperglutamylation (Pathak, Austin-Tse, Liu, Vasilyev, & Drummond, 2014) impact ciliary motility in different model organisms, revealing that precisely controlled levels of glutamylation are crucial to maintain proper ciliary function. Several studies also showed that glutamylation regulates IntraFlagellar Transport (IFT) dynamics, a bidirectional motility of ciliary constituents along axonemal microtubules required for assembly and maintenance (Scholey, 2003). Polyglutamylation positively regulates the IFT and microtubules motors (O’Hagan et al., 2011; Sirajuddin, Rice, & Vale, 2014) whereas defective polyglutamylation seems to impair anterograde IFT dynamics (Hong et al., 2018). Consequently, glutamylation is also important for ciliary signaling, as it impacts the localization of signaling molecules, such as Polycystins (He et al., 2018; O’Hagan et al., 2011), or Sonic Hedgehog components (Hong et al., 2018). Given the variety of ciliary functions associated with polyglutamylation, it is not surprising that mutations in enzymes generating or removing polyglutamylation are linked to ciliopathies. Notably, ciliary motility defects linked to glutamylation imbalance have been associated with male infertility, airway defects and dysfunction in ependymal cells in mice (Campbell et al., 2002; Giordano et al., 2019; Grau et al., 2013; Konno et al., 2016; Mullen, Eicher, & Sidman, 1976; Vogel, Hansen, Fontenot, & Read, 2010; Wu, Wei, & Morgan, 2017). Also, hyperglutamylation of microtubules in neuronal axons has been associated with neurodegeneration in mice and human (Magiera et al., 2018; Shashi et al., 2018).

The photoreceptor cell possesses the largest cilium of the human body (called the outer segment), characterized by the stack of hundreds of membrane discs along the axonemal microtubules. This axoneme is connected to the cell body via a thin microtubule-based region called the connecting cilium (Mercey et al., 2022). Mutations of the gene *AGBL5*, coding for the deglutamylase CCP5, have been associated with Retinitis pigmentosa, the most prevalent family of inherited retinal diseases leading to photoreceptor death (Astuti et al., 2016; Branham et al., 2016; Kastner et al., 2015). This provided the first link between tubulin polyglutamylation and photoreceptor degeneration in humans. Recently, a mouse model lacking CCP5 showed that hyperglutamylation observed in this mutant leads to photoreceptor cell degeneration with altered outer segments (Aljammal et al., 2024). However, mechanisms of photoreceptor cell degeneration at play are not known.

Here, we elucidate the role of tubulin PTMs in the outer segment (OS) of mouse photoreceptor cells. Using super resolution Ultrastructure expansion microscopy (U-ExM) technique allowed us to obtain a deeper understanding of tubulin PTMs distribution inside the OS with nanoscale precision. Analyzing mutant mice deficient for key enzymes of tubulin glutamylation, but also acetylation, we unraveled that imbalance of PTMs inside the OS leads to structural defects which impair photoreceptor function and can lead to retinal degeneration.

## Results

### The outer segment of mouse rod photoreceptor cells outer is enriched in tubulin PTMs

Using Ultrastructure Expansion Microscopy (U-ExM) that we recently adapted for retina imaging (Gambarotto et al., 2018; Mercey et al., 2022), we first assessed the localization of tubulin PTMs along the rod photoreceptor cilium, - the outer segment - in mouse retina with an expansion factor of 4.2X (**Supplementary Figure 1A, B**). We focused on the longitudinal and transversal views comprising the basal body, the connecting cilium, where many retinopathy-associated proteins localize (Bachmann-Gagescu & Neuhauss, 2019), and the bulge region that we recently described as a crucial compartment for membrane disc formation (Faber et al., 2023) and the distal axoneme. To map the distribution of different PTMs along the axoneme of the outer segment, we co-stained all microtubules using a mixture of anti α- and β-tubulin antibodies with a panoply of antibodies specific to tubulin PTMs: mono-glycylation (TAP952), acetylation (Acetylated tubulin), glutamylation- and poly-glutamylation (GT335 and PolyE) and detyrosination (Detyrosinated tubulin) (**Figure 1A-E, Supplementary Figure 1B**).

**Figure 1:**
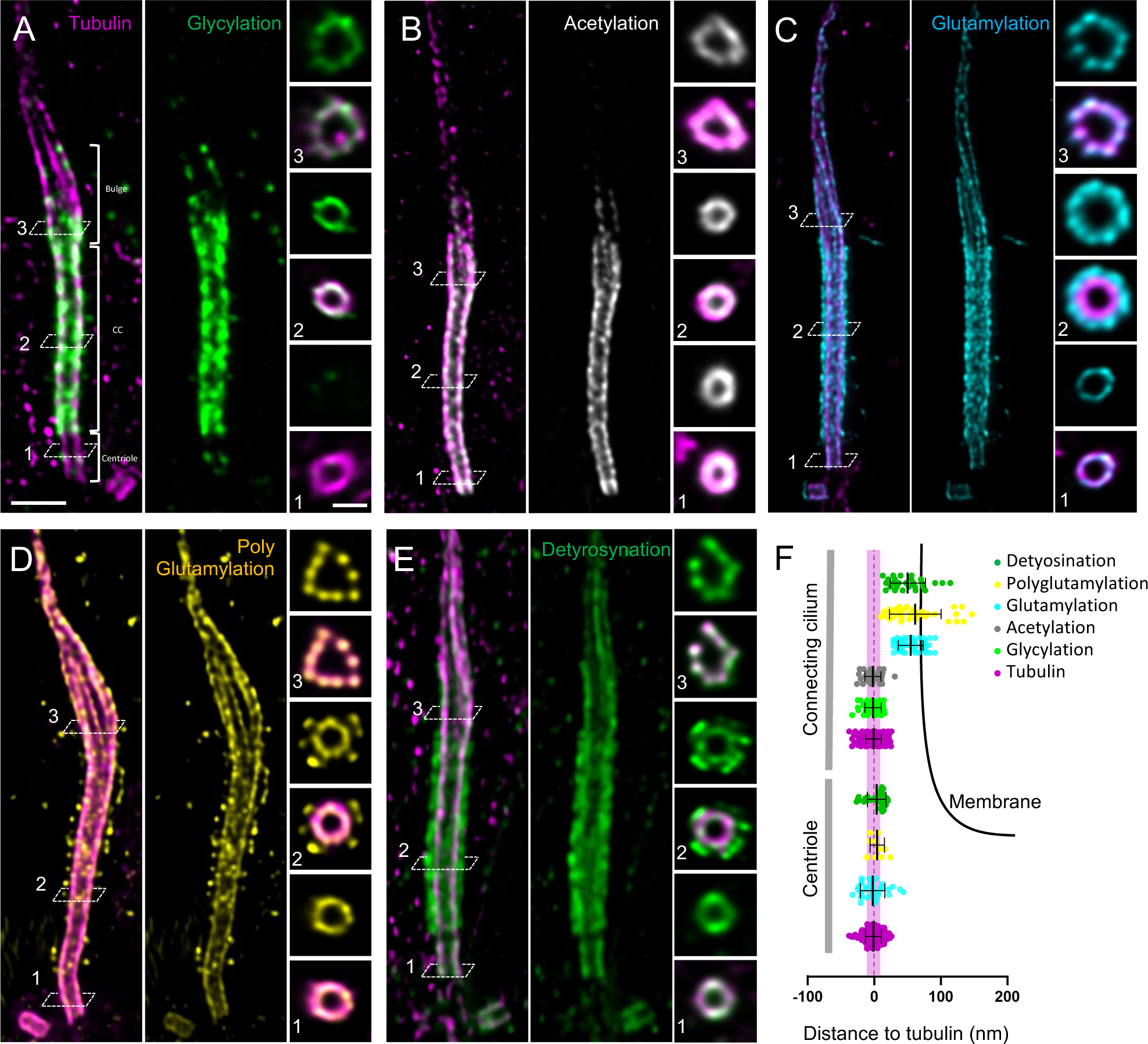
Molecular mapping of tubulin PTMs in mouse photoreceptor cells. (**A-E**) Confocal images of expanded photoreceptor cells stained with tubulin and (A) glycylation (TAP952, green), (B) acetylation Tubulin (gGray), (C) glutamylation (GT335, cyan), (D) polyglutamylation (PolyE, yellow) or (E) detyrosination (green). Scale bar: 500 nm. Transversal section images corresponding to different regions of the OS (centriole, connecting cilium and bulge depicted by the dashed lines and numbers on longitudinal images) are represented on the right side. Scalebar: 200 nm. (**F**) Schematic representation of the outer segment centriole and connecting cilium on which measured distances to tubulin of different proteins are represented. Membrane is depicted as a black line. Width of a microtubule (20nm) is depicted in magenta as a reference. Glutamylation (centriole): -1.21 nm +/- 18.61 (N=3); polyglutamylation (centriole): 5.66 nm +/- 10.91 (N=1); detyrosination (centriole): 4.92 nm +/- 14.07 (N=2); glycylation (CC): -0.57 nm +/- 12.45 (N=3); acetylation (CC): -0.93 nm +/- 12.30 (N=2); glutamylation (CC): 56.42 nm +/- 18.90 (N=3); polyglutamylation (CC): 63.5 nm +/- 39.01 (N=3); detyrosination (CC): 51.98 nm +/- 26.94 (Mean +/- SD). Tubulin is used as a reference : 0 nm +/- 12 (N>10).

We first mapped the tubulin glycylation. This modification has been mainly found in motile cilia/flagella (Gadadhar et al., 2021; Grau et al., 2013) but is also present in primary cilia assembly (Gadadhar et al., 2017; Rocha et al., 2014). In the photoreceptor cilium that can be considered as a specialized primary cilium, we found that glycylation is restricted to the connecting cilium and the bulge region, where it exactly lines microtubules (-0.6 nm shift relative to α- and β-tubulin staining) (**Figure 1A, F**). In contrast, glycylation was mostly absent distally to the bulge and we confirmed that it was also absent from the basal body (Guichard, Laporte, & Hamel, 2023). However, by oversaturating the signal, we found a reproducible faint signal localizing at the level of subdistal appendages of the centriole (**Supplementary Figure 2**).

We next analyzed tubulin acetylation status in the outer segment. This modification takes place in the lumen of the microtubules (**Supplementary Figure 1C**), acting on their mechanical properties (Eshun-Wilson et al., 2019). We found that acetylation is present from the basal body to the bulge region of the axonemal microtubules (**Figure 1B, F**). However, similar to glycylation, the signal above the bulge was mostly absent or faint, suggesting that distal disorganized axonemal microtubules are less prone to be modified by glycylating or acetylating enzymes.

Next, we examined glutamylation localization with two different antibodies: GT335 raised against a synthetic peptide mimicking the glutamylation modification on a C-terminal tubulin tail; and PolyE, an antibody recognizing long glutamate side chains (> 3 glutamates) (Rogowski et al., 2010; Shang, Li, & Gorovsky, 2002) (**Figure 1C, D**). For both antibodies, glutamylation staining reveals two localizations along the photoreceptor outer segment. The basal body is decorated with both glutamylation and poly-glutamylation with the exact same localization as the tubulin, as previously shown in *Chlamydomonas* and human centrioles (Gambarotto et al., 2019; Mahecic et al., 2020) (**Figure 1C, D, F**). However, the pattern changes drastically once the connecting cilium (CC) starts, with glutamylated tubulin (GT335 antibody) and polyglutamylated tubulin (polyE) signals more external to the microtubule wall and all along the CC, forming a sheath-like structure. We quantified this signal about 60 nm away from the microtubule center of mass (**Figure 1C, D, F**), rather close to the membrane, where CEP290 has been recently localized (Mercey et al., 2022) (**Figure 1F**). The discrepancy between the position of tubulin and that of glutamylation suggests that this modification might decorate another protein(s) than tubulin, which are located closer to the membrane. This external glutamylation signal is restricted to the connecting cilium, whereas the bulge and the distal axoneme exhibit glutamylation signal at the level of the microtubules, similar to the centriole. Moreover, unlike acetylation or glycylation signals, polyglutamylation seems to propagate more distally in the outer segment axoneme. To get a better impression of the precise localization, we also looked at the transverse view, where we confirmed the presence of a sheath-like glutamylation signal at the connecting cilium (**Figure 1C, D**). Additionally, polyglutamylation staining was observed on the tubulin, a signal not detected by the GT335 glutamylation antibody (**Figure 1D**). We also noticed that in some cases, PolyE accumulates at the base of the cilium, a signal that resembles IFT train accumulation in U-ExM (**Supplementary Figure 3**, white arrows) (Van den Hoek et al., 2022).

Finally, we also assessed the profiles of detyrosinated tubulin and delta2-tubulin. These two tubulin PTMs consist in the removal of the gene-encoded C-terminal tyrosine (detyr-tubulin), and the further cleavage of the penultimate glutamate residue (Δ2-tubulin) of α-tubulin (**Supplementary Figure 1C**). The staining pattern of detyrosinated-tubulin resembled what we have found for glutamylation, and in particular for polyglutamylation as shown in the transverse view at the level of the CC (**Figure 1E**). Detyrosinated tubulin is thus localized on microtubules, but also forms a sheath-like structure further away from them. Staining for Δ2-tubulin, by contrast, yielded a signal that was not distinct enough to conclude about the precise localization of this PTM (**Supplementary Figure 4**).

Altogether, these data reveal a complex distribution of tubulin PTMs along the photoreceptor outer segment. We found a specific enrichment at the level of the CC, where all PTMs analyzed are present, hinting at a prominent role of these modifications in this compartment.

### Perturbed tubulin PTMs lead to photoreceptor cell degeneration

Next, we investigated the physiological relevance of tubulin PTMs for photoreceptor cells. Indeed, it has been recently shown that mutation in *AGBL5*, coding the deglutamylase CCP5, lead to retinitis pigmentosa in humans. This suggests that perturbation of glutamylation could lead to drastic consequences on photoreceptor survival. We thus analyzed retina of mice lacking CCP5 (*Ccp5^-/-^*) and CCP1 (*Ccp1^-/-^*). Absence of these two deglutamylating enzymes is expected to lead to hyperglutamylation (Rogowski et al., 2010). To test whether other PTMs than glutamylation could affect photoreceptor cells, we also assessed the consequence of loss of tubulin acetylation in *Atat1^-/-^* mice.

After expansion, we assessed the integrity of the retina in mice aged from 8 to 18 Months old using rhodopsin, a marker of the rod outer segment, as well as α- and β-tubulin staining, in comparison to control mice (**Figure 2A-D**). Knockout of the deglutamylases CCP1 or CCP5 lead to a severe photoreceptor degeneration at 8 and 12 months, respectively (**Figure 2B, C**). This is highlighted by the substantial decrease of ONL thickness compared to WT, where only a couple of nuclear rows are remaining (**Figure 2B, C, E, Supplementary Figure 5A**). This phenotype was already described in pcd (purkinje cell degeneration) mice, bearing an inactivating mutation in the *AGTPBP1* gene (coding for CCP1) (LaVail, Blanks, & Mullen, 1982). Additionally, we show a mislocalization of rhodopsin in these mutants, with a signal at the ONL-surrounding nuclei; a hallmark of photoreceptor degeneration (**Figure 2B, C**, white arrows and insets). Interestingly, we noticed during dissections that retinas from *Ccp5^-/-^* mice are thinner and more fragile compared to WT (**Supplementary Figure 5B**). Of note, the other layers of the retina were not affected in these mutants. By contrast, we show that ATAT1 deficiency has no overall impact on photoreceptor survival, even in 18-month-old mice (**Figure 2D**). Indeed, the thickness of the ONL is not significantly changed compared to control, and the rhodopsin staining reveals the presence of organized outer segments (**Figure 2D, E**).

**Figure 2:**
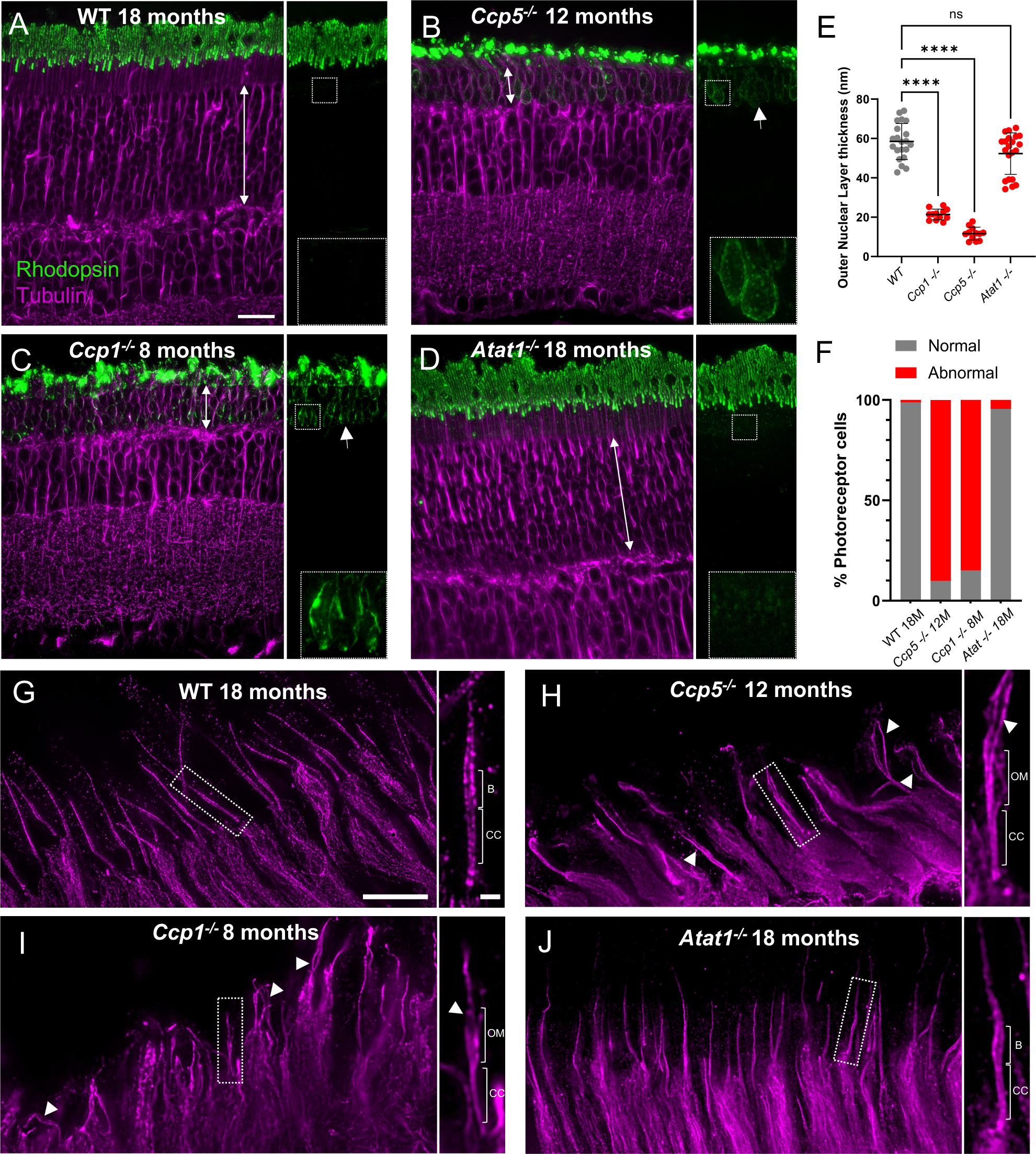
Deglutamylase mutants lead to retinal degeneration with photoreceptor axonemal disorganization. (**A-D**) Expanded retina at low magnification stained for tubulin (magenta) and rhodopsin (green) in WT (A), *Ccp5^-/-^* (B), *Ccp1^-/-^* (C) or *Atat1^-/-^* (D). On the right, the rhodopsin channel alone highlights the presence of the mislocalized signal in the ONL in the degenerating retina (white arrows and insets). Thickness of the Outer Nuclear Layer (ONL) is depicted with white double arrows. Scalebar: 20 µm. (**E**) Measurements of the ONL thickness in the different conditions, corrected by the expansion factor. WT: 58.47 nm +/- 9.15 (N=3); *Ccp1^-/-^*: 21.36 nm +/- 2.76 (N=2); *Ccp5^-/-^*: 11.65 nm +/- 3.27 (N=2); *Atat1^-/-^*: 52.31 nm +/- 10.54 (N=3) (Mean +/- SD) (**F**) Percentage of individual photoreceptor cells with normal or abnormal axonemal structures. WT: normal: 98.89%, abnormal: 1.11% (N=3); *Ccp5^-/-^*: normal: 9.84%, abnormal: 90.16% (N=2); *Ccp1^-/-^*: normal: 15.1%, abnormal: 84.9% (N=2); *Atat1^-/-^*: normal: 95.5%, abnormal: 4.5% (N=3) (**G-J**) Expanded photoreceptor cells stained with tubulin (magenta) highlighting the axonemal structure in WT (G), *Ccp5^-/-^* (H), *Ccp1^-/-^* (I) or *Atat1^-/-^* (J). Scalebar: 5 µm. Insets depicted with dashed lines are represented on the right. Scalebar: 500 nm. CC: Connecting Cilium; B: Bulge; OM: Open microtubules. White arrowheads show open microtubules in low magnification images.

We next focused the analysis on the photoreceptor outer segment, where PTMs are generally enriched (**Figure 2G-J**, **Figure 1A-E**). Given that CCP5 deficiency is linked to retinitis pigmentosa, and that we recently showed that FAM161A-associated retinitis pigmentosa RP28 is due to structural defects at the level of the CC, we investigated whether this structural element was affected in deglutamylase mutants.

Using tubulin staining to reveal the architecture of the ciliary axoneme, we showed that WT photoreceptor OS have straight axonemal microtubule extending toward the distal part of the cilium (**Figure 2G**). We also noticed the presence of the bulge region, delineating the end of the CC (**Figure 2G, Supplementary Figure 1A**). By contrast, in the two deglutamylase mutants analyzed, the structure of the OS is highly impaired, mostly with open axoneme on its distal end (**Figure 2H, I,** white arrowheads). Interestingly, we found that for *Ccp5^-/-^* and *Ccp1^-/-^* mice, the structural defects were observed mostly above the CC, whereas the tubulin shaft seemed preserved at the CC, as it has been described in mice mutant for the bulge protein LCA5 (Faber et al., 2023). The fact that more than 80% of the photoreceptor cells analyzed are defective in *Ccp5^-/-^* and *Ccp1^-/-^* mutants highlights the high penetrance of the degenerative phenotype in these mutants (**Figure 2F**). By comparison, *Atat1^-/-^* photoreceptor cells have no overall axonemal defects in old mice (**Figure 2F, J**), demonstrating that lack of tubulin acetylation does not result in obvious alteration at the level of photoreceptor cells.

### Distal axoneme disorganization in the outer segment and ciliary transport defects in deglutamylase deficiency

To gain mechanistic insights into photoreceptor degeneration linked to hyperglutamylation, and more specifically to CCP5 mutations which lead to retinitis pigmentosa in human, we next mapped the molecular composition of photoreceptors by U-ExM with a specific focus on the outer segment (OS). We analyzed staining for different PTMs (glycylation, glutamylation and acetylation), the CC marker POC5 (Mercey et al., 2022), the bulge marker LCA5 (Faber et al., 2023), and the Intraflagellar Transport (IFT) component IFT88.

Since lack of both CCP1 and CCP5 affect the glutamylation status, we first examined the GT335 pattern along the photoreceptor OS. As expected, we observed a strong hyperglutamylation in CCP5 mutants where the signal extends towards the distal part of the cilium, thus decorating the whole axoneme (**Figure 3A, G**). Remarkably, a similar pattern was observed for acetylated tubulin, where the signal was prolonged distally (**Figure 3B, H**). Low magnification images revealed that hyperglutamylation is not restricted to the OS but extends to the inner segment of the photoreceptor cells in CCP5 mutants (**Supplementary Figure 6**). Of note, in CCP1 mutants, axonemes appear shorter than in CCP5 mutants, probably explaining why hyperglutamylation and hyperacetylation towards the distal axoneme were less pronounced in these mutants. We also noticed that CCP5, but not CCP1 mutants display a strongly reduced glycylation signal along the axoneme, underpinning that hyperglutamylation caused by the loss of some of the deglutamylases could lead to reduced glycylation, as previously described (Grau et al., 2017) (**Figure 3C, I**).

**Figure 3:**
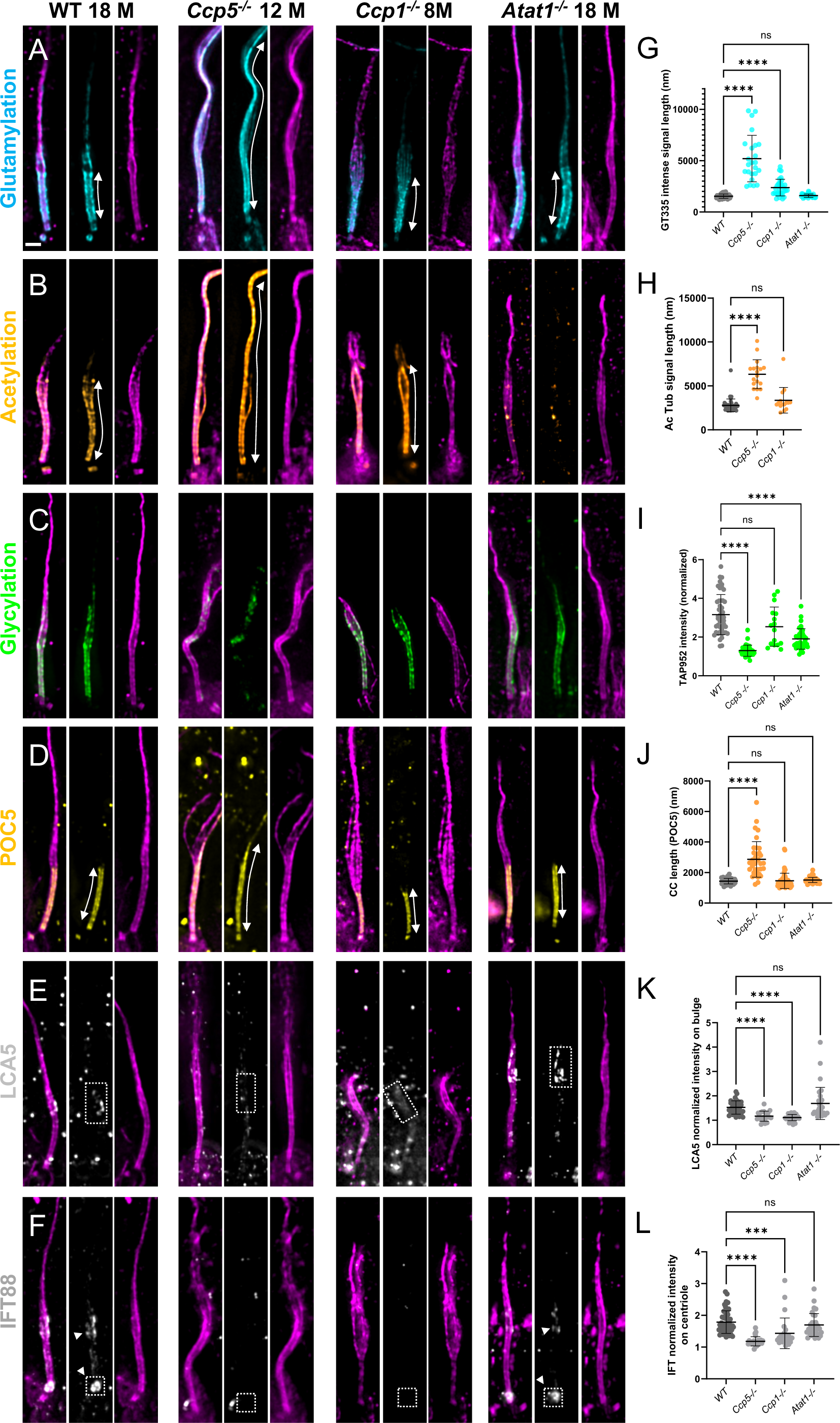
Deglutamylase mutants cause imbalance in PTMs paralleled with defects in outer segment structure and transport. (**A-F**) Expanded photoreceptor outer segments observed in WT, *Ccp5^-/-^*, *Ccp1^-/-^* or *Atat1^-/-^* and stained with GT335 (cyan) (A), Acetylated tubulin (orange) (B), TAP952 (green) (C), POC5 (yellow) (D), LCA5 (gray) (E), and IFT88 (gray) (F). White double arrows depict the length of the signal measured in (G), (H), (J). Rectangles with dashed lines shows the area where intensity measurements were picked in (K) and (L). White arrowheads show the dual localization of IFT at the bulge and at the cilium entry. Scalebar: 500 nm. (**G**) GT335 intense signal length corrected by the EF. WT: 1546 nm +/- 212.8; *Ccp5^-/-^*: 5209 nm +/- 2260; *Ccp1^-/-^*: 2378 nm +/- 814.2; *Atat1^-/-^*: 1594 nm +/- 166.1 (**H**). Acetylated tubulin signal length in the outer segment corrected by the EF. WT: 2795 nm +/- 729.4; *Ccp5^-/-^*: 6337 nm +/- 1645; *Ccp1^-/-^*: 3365 nm +/- 1463 (**I**) Intensity measurement of TAP952 signal along the outer segment CC normalized on background. WT: 3.16 nm +/- 1.03; *Ccp5^-/-^*: 1.30 +/- 0.29; *Ccp1^-/-^*: 2.54 +/- 1.01; *Atat1^-/-^*: 1.90 +/- 0.53 (**J**) CC length measured with POC5 and corrected by the EF. WT: 1433 nm +/- 179; *Ccp5^-/-^*: 2864 nm +/- 1164; *Ccp1^-/-^*: 1458 nm +/- 498; *Atat1^-/-^*: 1512 nm +/- 167 (**K**) LCA5 intensity at the bulge normalized on background. WT: 1.53 +/- 0.27; *Ccp5^-/-^*: 1.17 +/- 0.21; *Ccp1^-/-^*: 1.11 +/- 0.13; *Atat1^-/-^*: 1.69 +/- 0.66 (**L**) IFT88 intensity at the cilium base normalized on background. WT: 1.79 +/- 0.36; *Ccp5^-/-^*: 1.18 +/- 0.15; *Ccp1^-/-^*: 1.44 +/- 0.48; *Atat1^-/-^*: 1.70 +/- 0.36. (Mean +/- SD). Animals: WT: N=3; *Ccp5^-/-^:* N=2; *Ccp1^-/-^*: N=2; *Atat1^-/-^*: N=3.

In both CCP1 and CCP5 mutants, we confirmed a massive disorganization of the microtubules above the CC, with axoneme opening (**Figure 3A-F**). Inside the CC, POC5 signal remains unaffected, confirming that the cohesion of the microtubules is still effective in this compartment, and that the observed photoreceptor cell degeneration is not due to structural defects of the CC. However, CCP5 mutants exhibit an exacerbated CC, with the POC5 signal extending to more than 5 µm in some cases (compared to 1.5 µm in the control mice) (**Figure 3D, J**). We confirmed this result by staining with another marker of the CC, CEP290 (**Supplementary Figure 7**). In contrast, CCP1 deficiency did not affect the length of the POC5 signal.

Next, we analyzed the bulge region, by investigating the distribution of LCA5 (Faber et al., 2023). We found that the deficiency of deglutamylases CCP1 or CCP5 leads to a highly reduced and diffused signal of LCA5, suggesting that the bulge region is lost, thus preventing membrane disc formation and presumably causing photoreceptor death (**Figure 3E, K, Figure 2B, C**).

As CCP1 and CCP5 mutants display distal axoneme disorganization, rhodopsin mislocalization throughout the ONL, and lack of LCA5, we hypothesized that these phenotypes could be linked to intraflagellar transport (IFT) defects. Indeed, we recently showed that the bulge region, marked by LCA5, is crucial to organize IFT at the level of photoreceptor cell, and that the loss of the bulge is associated with defects in the IFT components localization (Faber et al., 2023). In WT, IFT88 accumulates both at the base of the cilium, where trains are formed, and above the CC, at the bulge (**Figure 3F**). By comparison, in both CCP1 and CCP5 mutants, the IFT88 signal is greatly reduced above the CC and the cilium entry, highlighting the defects in trafficking towards the photoreceptor outer segment (**Figure 3F, L**).

In parallel, we assessed the impact of alpha tubulin acetyltransferase 1 mutants, which undergo a complete signal loss of acetylation (**Supplementary Figure 8**). We first show that these mutant mice have no structural defects at the level of the outer segment (**Figure 3A-F**), confirming results obtained at the level of the whole retina (**Figure 2D**). Moreover, these mutants exhibit no obvious changes in the glutamylation status of the outer segment (**Figure 3A, G**). However, we noticed a reduced level of glycylation in *Atat1^-/-^* mutants (**Figure 3C, I**). The staining of the CC marker - POC5 - revealed no difference in the CC length compared to WT (**Figure 3D, J**), suggesting that this structure is intact. In line with this, the bulge marker LCA5 was not impacted by the loss of acetylation in the OS (**Figure 3E, K**). Finally, in *Atat1^-/-^* mutants, we show that the dual localization of IFT88 is conserved (**Figure 3F**, white arrowheads) and that the intensity of IFT88 staining is similar to the WT at the base of the cilium (**Figure 3L**).

Altogether, we demonstrate that defects in glutamylation, but not acetylation, strongly impact the architecture of the OS axoneme, that is associated with transport defects.

### Progressive disorganization of photoreceptor OS in CCP5 mutant mice

To better define the time course of the CCP5-associated retinal degeneration, we analyzed the anatomy of photoreceptor cilia in 3-, 7-, 10- and 12-month-old *Ccp5^-/-^* mice. Using rhodopsin and tubulin staining as readouts for retina integrity, we found that degeneration starts at around 3 months, when ONL is already about 10 µm thinner than in the WT (**Figure 4A, E**). ONL thickness progressively decreases in the following months to ultimately reach only 2 or 3 cell layers at 12 months (**Figure 4A-E**, white double arrows), highlighting a progressive degeneration over the first year after birth. This continuous decrease of the ONL thickness is accompanied by mislocalization of rhodopsin around nuclei, a defect that becomes obvious at 7 months (**Figure 4A-D**, white arrows). At the OS level, rhodopsin and tubulin signals reveal the overall absence of defects at 3 and 7 months, where membrane discs appear correctly organized, the bulge region is still present, and the distal axoneme is straight and properly organized (**Figure 4A, B**). However, from 10 months onwards, we observe a more and more pronounced disorganization of the OS, with opening of axonemes, shorter membrane stacks, and the presence of isolated rhodopsin patches outside the cell (**Figure 4C, D**, white arrowheads).

**Figure 4:**
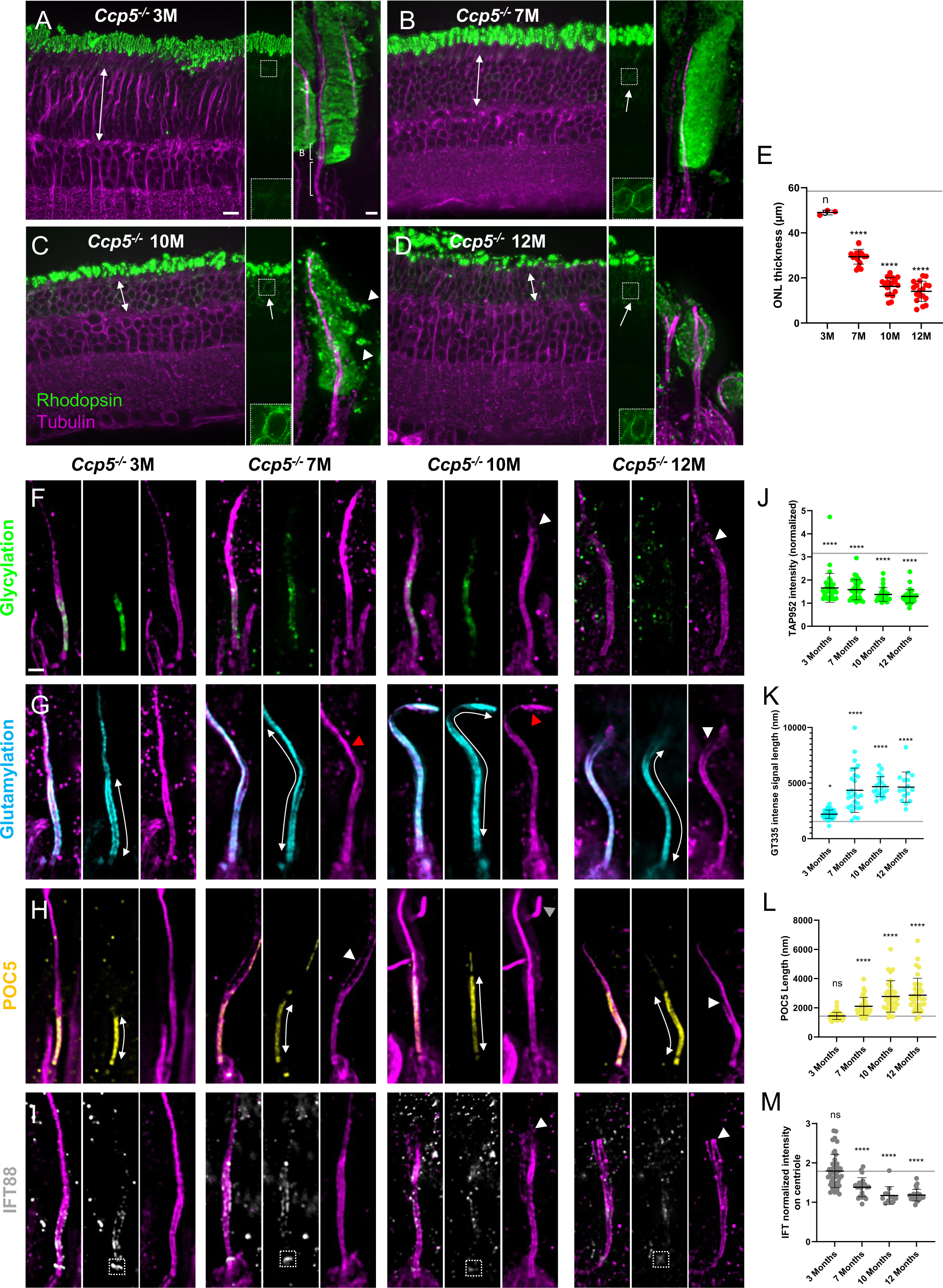
Dynamic of photoreceptor cell degeneration in CCP5 -/- mice. (**A-D**) Overview of photoreceptor cell progressive loss in expanded retina of 3 months- (A), 7 months- (B), 10 months- (C), 12 months- (D) old *Ccp5^-/-^* mice stained with rhodopsin (green) and tubulin (magenta). Scalebar: 10 µm. For each time point, a representative image of the rhodopsin channel alone is shown in the middle with an inset and a single photoreceptor cell is depicted on the right. White double arrows reveal the thickness of the ONL measured in (E). White arrows show the rhodopsin signal at the level of the nuclei. White arrowheads show floating rhodopsin signal outside of the outer segment. Scalebar: 500 nm. CC: connecting cilium, B: Bulge. (**E**) ONL thickness measurement and compared to WT (gray line). *Ccp5^-/-^* 3M: 49.0 nm +/- 1.1; *Ccp5^-/-^* 7M: 29.4 nm +/- 3.3; *Ccp5^-/-^* 10M: 16.2 nm +/- 3.9; *Ccp5^-/-^* 12M: 14.1 nm +/- 4.5 (Mean +/-SD). (**F-I**) Single cell expanded photoreceptor cells of 3 months, 7 months-, 10 months-, 12 months-old *Ccp5^-/-^* mice revealing progressive defects of the outer segment using TAP952 (green) (F), GT335 (cyan) (G), POC5 (yellow) (H) and IFT88 (gray) (I). White double arrows depict the length of the signal measured for GT335 and POC5. Red and white arrowheads point to curled and broken axonemes, respectively. Rectangles with dashed lines shows the area where intensity measurements were picked with IFT88 signal. Scalebar: 500 nm. (**J**) Intensity measurement of TAP952 signal along the outer segment CC normalized on background. *Ccp5^-/-^* 3M: 1.66 +/- 0.63; *Ccp5^-/-^* 7M: 1.58 +/- 0.43; *Ccp5^-/-^* 10M: 1.38 +/- 0.30; *Ccp5^-/-^* 12M: 1.30 +/- 0.29 (Mean +/-SD). (**K**) GT335 intense signal length corrected by the EF. *Ccp5^-/-^* 3M: 2206 nm +/- 374; *Ccp5^-/-^* 7M: 4355 nm +/- 1990; *Ccp5^-/-^* 10M: 4678 nm +/- 903; *Ccp5^-/-^* 12M: 4640 nm +/- 1357 (Mean +/-SD). (**L**) CC length measured with POC5 and corrected by the EF. *Ccp5^-/-^* 3M: 1446.0 nm +/- 241; *Ccp5^-/-^* 7M: 2100 nm +/- 609; *Ccp5^-/-^* 10M: 2779 nm +/- 1076; *Ccp5^-/-^*12M: 2864 nm +/- 1164 (Mean +/-SD). (**M**) IFT88 intensity at the ciliary base normalized on background. *Ccp5^-/-^*3M: 1.80 +/- 0.42; *Ccp5^-/-^* 7M: 1.36 +/- 0.24; *Ccp5^-/-^*10M: 1.17 +/- 0.23; *Ccp5^-/-^* 12M: 1.18 +/- 0.15 (Mean +/-SD). For each measurement, WT baseline is depicted with a gray line. Measurements were compared to WT values obtained figure 2 or 3, and 12 month-old measurements are the same as in figure 2 or 3. Animals: 3 months-, 7 months-, 10 months-, 12 months-old *Ccp5^-/-^* mice: N=2.

We next analyzed several markers of the OS during the time course of degeneration to obtain insights into the molecular mechanisms involved in CCP5-associated retinitis pigmentosa. At 3 months, we find that glycylation staining is strongly reduced concomitantly with highly increased glutamylation (**Figure 4F, G, J, K**). By contrast, at this timepoint, both the CC length (POC5) (**Figure 4H, L**) and the IFT enrichment at the basal body (IFT88) (**Figure 4I, M**) are comparable to the WT, suggesting that the PTM imbalance precedes the structural and functional defects of the OS. From 7 months onwards and gradually increasing up to 12 months, we find that axoneme disorganization is more and more pronounced, with distal axonemal microtubules mostly curled or broken (**Figure 4A-I**, red and white arrowheads, respectively). TAP952 (glycylation) and IFT88 (Intraflagellar Transport) intensities are progressively reduced and ultimately absent from 12-month-old photoreceptor cells (**Figure 4F, I, J, M**). In parallel, the length of the CC gradually increases (**Figure 4H, L**) alongside with an increase in hyperglutamylation that decorates the whole distal axoneme from 10 months of age (**Figure 4G, K**).

Altogether, we showed that CCP5 mutants exhibit a slow and progressive degeneration of the photoreceptor cells characterized by the progressive decrease of glycylation and a concomitant increase of hyperglutamylation. At later stages, the signal of ciliary transport proteins is progressively reduced, in parallel with a disorganization of the axonemal microtubules. The loss of the bulge region accompanied by the exacerbation of the CC is presumably leading to the inability to form new membrane discs, ultimately causing photoreceptor cell death.

### Molecular mapping of the tubulin PTMs in human photoreceptor cell outer segment

Finally, we wanted to assess whether the observed distribution of tubulin PTMs is conserved in human retina, as mutations of *AGBL5*, coding for CCP5, leads to retinitis pigmentosa (**Figure 5A-F**). To do so, we expanded human retina from a healthy adult. We first noticed that the length of the CC is shorter in human photoreceptor cells as compared to mouse. Using either GT335 or POC5 (Mercey et al., 2022) as markers of the CC, we demonstrate that the human CC is less than half the length of the one in mouse, spanning about 650 nm vs approximatively 1600 nm in mouse (**Figure 5G, Supplementary Figure 9A**). Interestingly, we also noticed that cone photoreceptors exhibit particularly long daughter centrioles that are more than 700 nm long, which could represent one of the biggest centrioles found in the human body (**Supplementary Figure 9B**).

**Figure 5:**
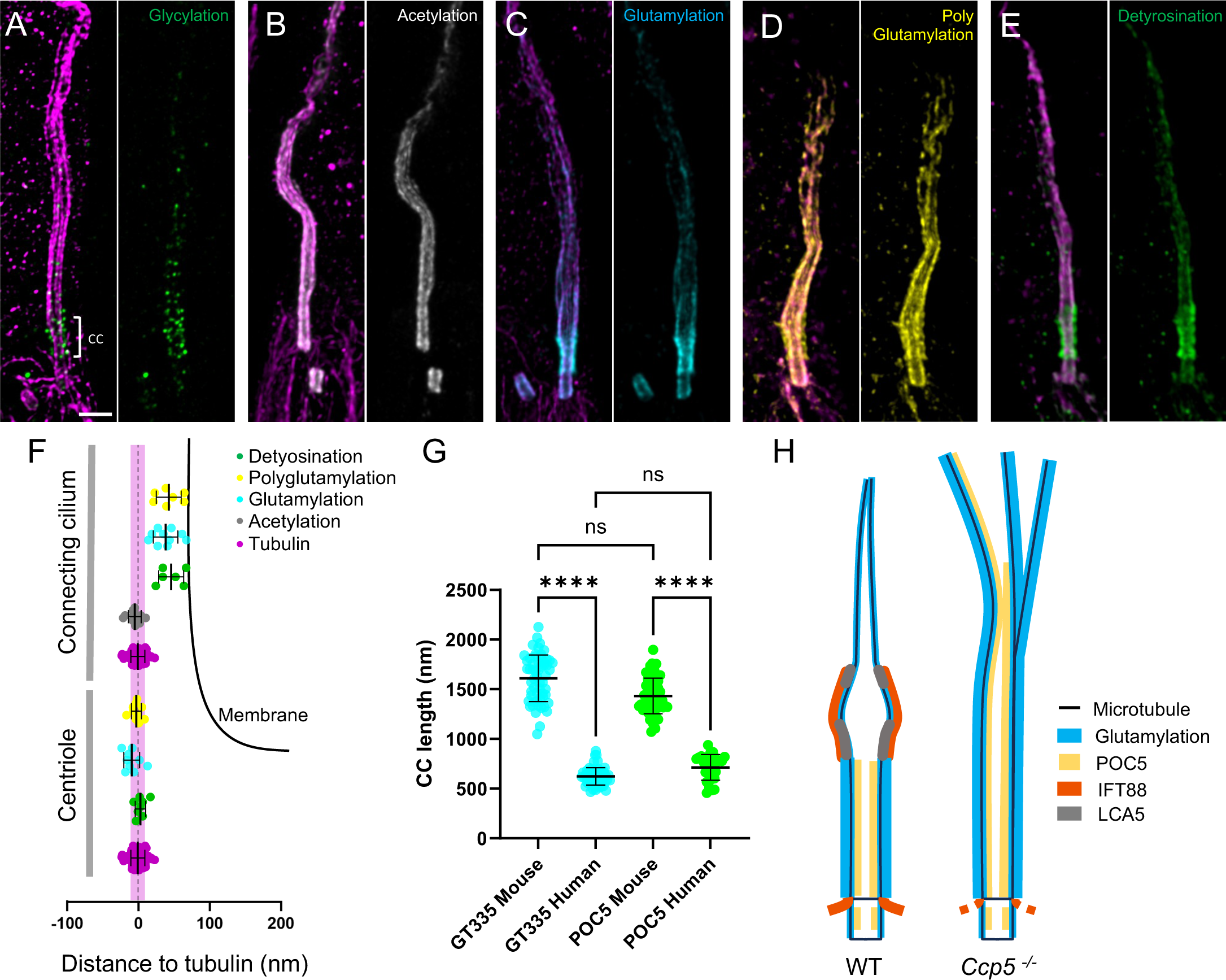
Molecular mapping of tubulin PTMs in human photoreceptor cells. (**A-E**) Confocal images of expanded photoreceptor cells stained for tubulin (magenta) and (A) glycylation (TAP952, green), (B) acetylation (gray), (C) glutamylation (GT335, cyan), (D) polyglutamylation (PolyE, yellow) or (E) detyrosination tubulin (green). Scale bar: 500 nm. (**F**) Schematic representation of the outer segment centriole and connecting cilium on which measured distances to tubulin of different proteins are represented. Detyrosination (centriole): 3.73 nm +/- 7.29; glutamylation (centriole): -8.60 nm +/- 11.16; polyglutamylation (centriole): -2.03 nm +/- 7.12; acetylation (CC): -4.41 nm +/- 9.03; detyrosination (CC): 47.15 nm +/- 17.70 glutamylation (CC): 39.25 nm +/- 17.44; polyglutamylation (CC): 43.78 nm +/- 17.68; (Mean +/- SD). Membrane is depicted as a black line. Width of a microtubule (20nm) is depicted in magenta as a reference. (**G**) Comparison of mouse and human CC length using GT335 and POC5 as markers. GT335 (mouse): 1610 nm +/- 235; GT335 (human): 623 nm +/- 87; POC5 (mouse): 1433 nm +/- 179; POC5 (human): 714 nm +/- 130 (Mean +/- SD). Sample replicate: N=1. (**H**) Model summarizing the defects observed in a *Ccp5^-/-^*photoreceptor cell Outer Segment compared to WT. Loss of CCP5 leads to hyperglutamylation (cyan) that propagates towards the distal part of the cilium. This is accompanied by the loss of the bulge region delineated by LCA5 (gray), the elongation of the CC marked with POC5 (yellow), and IFT defects (red). Consequently, the distal part of the axoneme is disorganized.

We show that glycylation signal (TAP952), while being faint, is present along the human CC and probably on the bulge, similarly to what we described in mouse (**Figure 5A**). However, the signal was not strong enough to allow quantifications. We also found that acetylation is decorating microtubules all along the OS, from the basal body to the bulge, but also distally, which was less obvious in mouse (**Figure 5B, F**). Next, analyzing glutamylation (GT335 and PolyE) and detyrosination signals, we observed the same pattern as in mouse, forming a sleeve outside of the microtubules at the level of the CC, and lining the microtubules at the basal body and the cilium above the CC (**Figure 5C-E**). The measured diameters revealed the same range of distances to microtubules as in mouse (**Figure 5F**, **Figure 1F**), highlighting the conservation of PTM localization along the photoreceptor outer segment in mouse and human. In summary, U-ExM unveiled the conservation of PTM distribution along the photoreceptor cell outer segment but also highlighted structural difference between mouse and human photoreceptors cells.

## Discussion

The photoreceptor outer segment, reaching a length of 50 µm in the human eye, is organized around its microtubule-based axoneme, and provides structural support for the regularly stacked membrane discs. As we recently described, disorganization of the axonemal structure leads to massive outer segment collapse, causing photoreceptor death (Mercey et al., 2022). Therefore, cellular determinants assuring the integrity of axonemal microtubules are expected to be of prime importance for the function and survival of photoreceptor cells. Besides structural features such as the inner scaffold inside the connecting cilium that directly maintains axoneme cohesion by connecting neighboring microtubule doublets (Mercey et al., 2022), tubulin PTMs have emerged as molecular actors of microtubule stability and function. The importance of tubulin PTMs in photoreceptor maintenance has recently been highlighted by the fact that mutations of *AGBL5*, coding the deglutamylase CCP5, lead to retinitis pigmentosa in human. However, mechanisms of photoreceptor degeneration associated with CCP5 deficiency have remained unknown.

Here, we explored the molecular localization of PTMs and assessed consequences of PTMs perturbations for the photoreceptor outer segment. We first provided a molecular mapping of 4 different tubulin PTMs: glycylation, acetylation, glutamylation (mono- and poly-) and detyrosination and revealed that they form distinct patterns along the outer segment. First, glycylation is the only modification that is mostly absent from the basal body, besides a faint signal that could correspond to the subdistal appendages. Whether this modification is of interest to position or to build this crucial structure of the centriole remains to be elucidated. At the level of the connecting cilium, all the analyzed PTMs are present, with different localizations. Whereas acetylation and glycylation are observed on the microtubules, as expected, we were surprised to see that glutamylation and detyrosination exhibit a strong signal, restricted to the CC, but about 60 nm away from the microtubule signal center of mass.

This distance and the absence of tubulin staining at this location excluded the possibility that we detect microtubule or tubulin trafficking along the CC. Therefore, glutamylation and detyrosination might decorate other substrates, particularly enriched along the inner part of the CC membrane. It has been recently shown that RPGR, a CC protein, is glutamylated by TTLL5 and is recognized by the GT335 antibody (Sun et al., 2016). Another possible target could be a component of the Y-links structures, that connect microtubule doublets to the membrane. We recently showed that CEP290 localizes at the level of the Y-links of photoreceptor cells and colocalizes with GT335 signal (Louvel et al., 2023; Mercey et al., 2022). Since CEP290 Carboxy terminal (Cter) sequence is enriched in glutamates, this protein could be a substrate of glutamylation, and therefore be recognized by glutamylation antibodies. However, the antibody against detyrosinated tubulin is supposed to recognize specifically the C-terminal sequence of alpha-tubulin. Why only the CC reveals such a pattern of detyrosination remains unknown, but we cannot exclude nonspecific signal. Interestingly, acetylation and glycylation seem almost totally absent above the bulge region, where axonemal microtubules are losing organization and even sometimes can be observed as singlets. One could imagine that glycylases and the tubulin acetyl transferase (ATAT1) localization is restricted to the region below the distal axoneme, where they act on the mechanical properties of axonemal microtubules, protecting them from a loss of cohesion. Localizing the enzymes responsible for PTMs along the outer segment would help to answer the different patterns observed.

We next analyzed the effect of PTM perturbation on photoreceptor cell maintenance, focusing on glutamylation, since mutations of the deglutamylase CCP5 lead to retinitis pigmentosa in human. Loss of either CCP1 or CCP5 deglutamylases leads to retinal degeneration in few months, where only 2 or 3 layers of photoreceptor nuclei remain at about one year of age (vs about 10 layers in WT, **Fig Supplementary 5A**). We used U-ExM to describe the degeneration process at the cellular level. Interestingly, the first obvious phenotype observed in these two mutant mice is the disorganization of the axoneme, that occurs above the CC (**Figure 5H**). This is distinct from what we described for *Fam161a* mutation (also leading to retinitis pigmentosa), where microtubules spread just above the basal body (Mercey et al., 2022), thus indicating a different molecular mechanism. The fact that the CC is mostly preserved from microtubule collapse in CCP1- and CCP5-deficient mice is highly similar to what we previously observed for the deficiency of the bulge protein Lebercilin (LCA5), causing Leber Congenital Amaurosis in human (Faber et al., 2023). Intriguingly, in Lebercilin-deficient mice, the bulge is no longer present, and CC markers exhibit longer signals, suggesting that Lebercilin could act as a ruler to dictate CC length. In CCP5 mutant mice, CC size is even more exacerbated, and LCA5 is no longer present, suggesting similar mechanisms in these two mutants, even if the onset of the degeneration is faster in LCA5 deficient mice (within the first month after birth) (**Figure 5H**).

We further showed that CCP5 loss leads to an important hyperglutamylation that is paralleled with the loss of glycylation, highlighting the competition between these two PTMs, as previously described for mice lacking glycylation in the retina (Grau et al., 2017) (**Figure 5H**). Importantly, hyperglutamylation is not restricted to the outer segment in CCP5 mutants, as the whole inner segment exhibits a strongly increased GT335 signal. We cannot exclude that hyperglutamylation of cytosolic microtubules in the cell body is partly responsible for photoreceptor cell death. Interestingly, defects of TTLL5, a glutamylase, leading to hypoglutamylation on its substrates is also leading to photoreceptor degeneration (Sun et al., 2016), showing that the fine-tuning of glutamylation is crucial to maintain the correct function of photoreceptor cells.

Surprisingly, direct comparison of CCP1- and CCP5-deficient mice at late stage revealed distinct effects on CC and on hyperglutamylation level. One reason for this might be the time course of degeneration between CCP1- and CCP5-deficient mice. CCP1 degeneration seems faster compared to CCP5, as the ONL thickness is comparable between 8-month-old *Ccp1*^-/-^ mice and 12-month-old *Ccp5*^-/-^ mice (**Figure 2E**). Moreover, *Ccp1*^-/-^ photoreceptor outer segments are shorter at 8-months compared to CCP5 mutants, reflecting a more advanced degeneration, where only a small portion of the axoneme is remaining. This would explain the difference in POC5 and GT335 signal length between *Ccp1*^-/-^ *and Ccp5*^-/-^ outer segments.

It has been shown for various types of cilia that impaired glutamylation leads to ultrastructural defects of the B-tubule, possibly impairing intraflagellar transport (IFT) (Yang et al., 2021). In photoreceptor cells, we observed a loss of IFT88 signal in both CCP1 and CCP5 mutant mice at the base of the cilium, similarly to what we previously showed in *Lca5* mutant mice (Faber et al., 2023) (**Figure 5H**). Interestingly, the loss of IFT recruitment in CCP5 mutant is concomitant to the disorganization of membrane discs of the outer segment in 10-month-old mice (**Figure 4C, I**). This result suggests that hyperglutamylation impairs IFT transport towards the bulge region, where membrane discs form, leading to the progressive collapse of the outer segment. One could imagine that hyperglutamylation change the IFT motor behavior. However, looking at the time course of photoreceptor degeneration in *Ccp5* mutant, hyperglutamylation of the axoneme happens before IFT88 loss, suggesting that perturbation of IFT is rather an indirect consequence of hyperglutamylation. We previously showed that IFT components are enriched at the bulge region, concentrating building blocks to form membrane discs mice (Faber et al., 2023). Since LCA5 signal at the bulge is also lost in deglutamylase mutants, a possible explanation is that hyperglutamylation leads to the loss of the bulge region, thus causing IFT components to diffuse to the distal cilium and preventing their recycling. Interestingly, we showed that in LCA5 mutant mice, glutamylation signal is also seen along the distal axoneme, suggesting that in physiological conditions, LCA5 could prevent hyperglutamylation at the level of the bulge (**Supplementary Figure 10**). How bulge-region-associated LCA5 and glutamylation are interconnected remains to be elucidated.

Finally, an open question is why the CC remains mostly intact in these mutants? How is this structure, particularly enriched in PTMs, maintained if transport is abolished? First, we previously showed that the inner scaffold maintains the cohesion of the axoneme by bridging neighboring microtubule doublets along the CC (Mercey et al., 2022). Second, it has been shown that in cilia with reduced glutamylation (resulting from the deficiency of TTLL5, TTLL6 or from overexpression of CCP5), a residual glutamylation signal is preserved at the level of the transition zone, suggesting that other enzymes could also regulate TZ glutamylation (He et al., 2018; Hong et al., 2018). In the photoreceptor outer segment, where the CC represents a 2-µm long TZ, the presence of additional enzymes could explain why this compartment is preserved from structural defects.

Our demonstration that tubulin PTMs are similarly present and distributed in human retina strongly suggests that observations made in the mouse models are relevant for human. One important difference we observed is the length of the connecting cilia, which in human is half the length of the mouse. This seems counterintuitive given that the outer segment of photoreceptors is longer in human as compared to mouse. A careful analysis in several species would help to understand how the length of the CC is regulated.

Altogether, our study revealed the importance of controlled levels of glutamylation in the highly specialized primary cilium of photoreceptor cells, the outer segment, to maintain the integrity of the axonemal structure. In addition, this work highlights the need to elucidate the subcellular events involved in retinal diseases such as retinitis pigmentosa. Indeed, this pathology being associated with mutations in about 80 genes, it is crucial to understand specific molecular mechanisms linked to each gene, to properly adapt therapeutic options to cure or slow down this type of diseases.

## Methods

### Mutant Mouse models

Animal care and use for this study were performed in accordance with the recommendations of the European Community (2010/63/UE) for the care and use of laboratory animals. Experimental procedures were specifically approved by the ethics committee of the Institut Curie CEEA-IC #118 (APAFIS #37315-2022051117455434 v2) in compliance with the international guidelines.

Mice used in this study have been described before: *Ccp1^-/-^* (PMID: 29449678), *Ccp5^-/-^*(pmid: 30635446), *Atat1^-/-^* (PMID: 23748901).

### Human tissue

The use of human samples was approved by the local Ethics Committee (CER-VD protocol No 340-15) and the patients signed informed consent.

### Ultrastructure Expansion Microscopy (U-ExM) of mouse and human retinas

Mice of desired age and genotype were sacrificed by cervical dislocation and their eyes were immediately collected and immersed in 4% PFA in PBS (n°15714S, Electron Microscopy Sciences). They were incubated overnight at 4°C, washed 3 x 20 min in PBS at room temperature and then stored at 4°C in PBS until required.

Human retina tissue was taken from a healthy retinal region of enucleated eyes due to tumor exenteration. Retina sample was fixed for 60 min in 4% PFA at RT and then wash in PBS before preparation for U-ExM.

Retina were then processed as described elsewhere (Faber et al., 2023; Mercey et al., 2022). Briefly, once flattened, retinas were placed inside the well of a 35 mm Petri dish (P35G-1.5-10-C, MatTek) for U-ExM processing. Retinas were first incubated overnight (ON) in 100 μL of 2% acrylamide (AA; A4058, MilliporeSigma) + 1.4% formaldehyde (FA; F8775, MilliporeSigma) at 37°C. The next day, solution is removed and 35 μL monomer solution composed of 25 μL of sodium acrylate (stock solution at 38% [w/w] diluted with nuclease-free water, 408220, MilliporeSigma), 12.5 μL of AA, 2.5 μL of N,N′-methylenebisacrylamide (BIS, 2%, M1533, MilliporeSigma), and 5 μL of 10× PBS was added for 90 minutes at RT. Then, MS was removed and 90 μL of MS was added together with ammonium persulfate (APS, 17874, Thermo Fisher Scientific) and tetramethylethylenediamine (TEMED, 17919, Thermo Fisher Scientific) as a final concentration of 0.5% for 45 min at 4°C first followed by 3 h incubation at 37°C to allow gelation. A 24-mm coverslip was added on top to close the chamber. Next, the coverslip was removed and 1 ml of denaturation buffer (200 mM SDS, 200 mM NaCl, 50 mM Tris Base in water (pH 9)) was added into the MatTek dish for 15 min at RT with shaking. Then, careful detachment of the gel from the dish with a spatula was performed, and the gel was incubated in 1.5 ml tube filled with denaturation buffer for 1 h at 95°C and then ON at RT. The day after, the gel was cut around the retina that is still visible at this step and expanded in 3 successive ddH2O baths. Then, the gel was manually sliced with a razorblade to obtain approximately 0.5 mm thick transversal sections of the retina that were then processed for immunostaining.

### Immunostainings

Gel slices were first incubated in 3 successive PBS 1X baths of 5 min. Then, gels were incubated with primary antibodies (Table S1) in PBS with 2% of Bovine Serum Albumin (BSA) overnight at 4°C. Gels were then washed 3 times 5 min in PBS with 0.1% Tween 20 (PBST) prior to secondary antibodies incubation for 3 h at 37°C. After a second round of washing (3 times 5 min in PBST), gels were expanded with three 10-min baths of ddH20 before imaging. Image acquisition was performed on an inverted confocal Leica Stellaris 8 microscope or on a Leica Thunder DMi8 microscope using a 20× (0.40 NA) or 63× (1.4 NA) oil objective with Lightning or Thunder SVCC (small volume computational clearing) mode at max resolution, adaptive as “Strategy” and water as “Mounting medium” to generate deconvolved images. 3D stacks were acquired with 0.12 μm z-intervals and an x, y pixel size of 35 nm.

### Quantifications

#### Expansion Factor

The expansion factor was calculated in a semiautomated way by comparing the full width at half maximum (FWHM) of photoreceptor mother centriole proximal tubulin signal with the proximal tubulin signal of expanded human U2OS cell centrioles using PickCentrioleDim plugin described elsewhere (Le Borgne et al., 2022). Briefly, more than 50 photoreceptor mother centrioles FWHM were measured and compared to a pre-assessed value of U2OS centriole width (25 centrioles: mean = 231.3 nm +/− 15.6 nm). The ratio between measured FWHM and known centriole width gave the expansion factor (**Supplementary Figure 1C**)

#### ONL thickness

ONL thickness was measured manually using tubulin staining on at least 2 different 20× original magnification images per replicate. Three measurements were performed per image to avoid bias due to retina dissection or slicing. Each measurement was subsequently corrected for the expansion factor.

#### Protein diameter

Using ImageJ, a line crossing centriole or connecting cilia on their diameter was drawn and plot profiles of each channel (protein of interest and tubulin) were generated. Then, distances between peak intensities of each protein were recorded. Average tubulin diameter was set at 170 nm from cryo ET data (Robichaux et al., 2019), to generate an expansion factor value for each protein measurement.

#### POC5, GT335 or Acetylated tubulin length

Protein signal lengths were measured using a segmented line drawn by hand (FIJI) to fit with photoreceptor curvature and corrected with the expansion factor.

#### Intensity measurements

Fluorescence intensity measurements of TAP952, LCA5 and IFT88 were performed on maximal projections using FIJI (46) on denoised images. The same rectangular region of interest (ROI) drawn by hand was used to measure the mean gray value of the protein signal and the corresponding background. Fluorescence intensity was finally calculated by dividing the mean gray value of the fluorescence signal by the mean gray value of the background (normalized mean gray value). For TAP952, measurements were performed all along photoreceptor CC and bulge, defined by tubulin. For LCA5 and IFT88, measurements were performed on the bulge region, and the basal body, respectively, defined by tubulin.

#### Statistics

The comparisons of more than 2 groups were made using nonparametric Kruskal–Wallis test followed by post hoc test (Dunn’s for multiple comparisons) to identify all the significant group differences. Every measurement was performed on at least 2 different animals, unless specified. Data are all represented as a scatter dot plot with centerline as mean, except for percentages quantifications, which are represented as histogram bars. The graphs with error bars indicate SD (+/−) and the significance level is denoted as usual (*p < 0.05, **p < 0.01, ***p < 0.001, ****p < 0.0001). All the statistical analyses were performed using Prism9. When possible, a minimum of 10 measurements has been performed per animal.

## Acknowledgements

This work was funded by the Swiss National Science Foundation (SNSF) grants 310030_205087 (VH, PG), the ProVisu and Gelbert foundations (VH, CK). CJ is supported by the French National Research Agency (ANR) awards ANR-20-CE13-0011, ANR-21-CE14-0045, and the Fondation pour la Recherche Medicale (FRM) grant MND202003011485. We thank the BioImaging Center of the University of Geneva, as well as K. Belloul, V. Dangles-Marie, V. Henriot, C. Jouhanneau (Institut Curie) for technical assistance. We thank Ronald Roepman for the use of *Lca5^gt/gt^* mouse.

## Data availability

No data was used for the research described in the article.

---

**Supplementary Figure 1:**
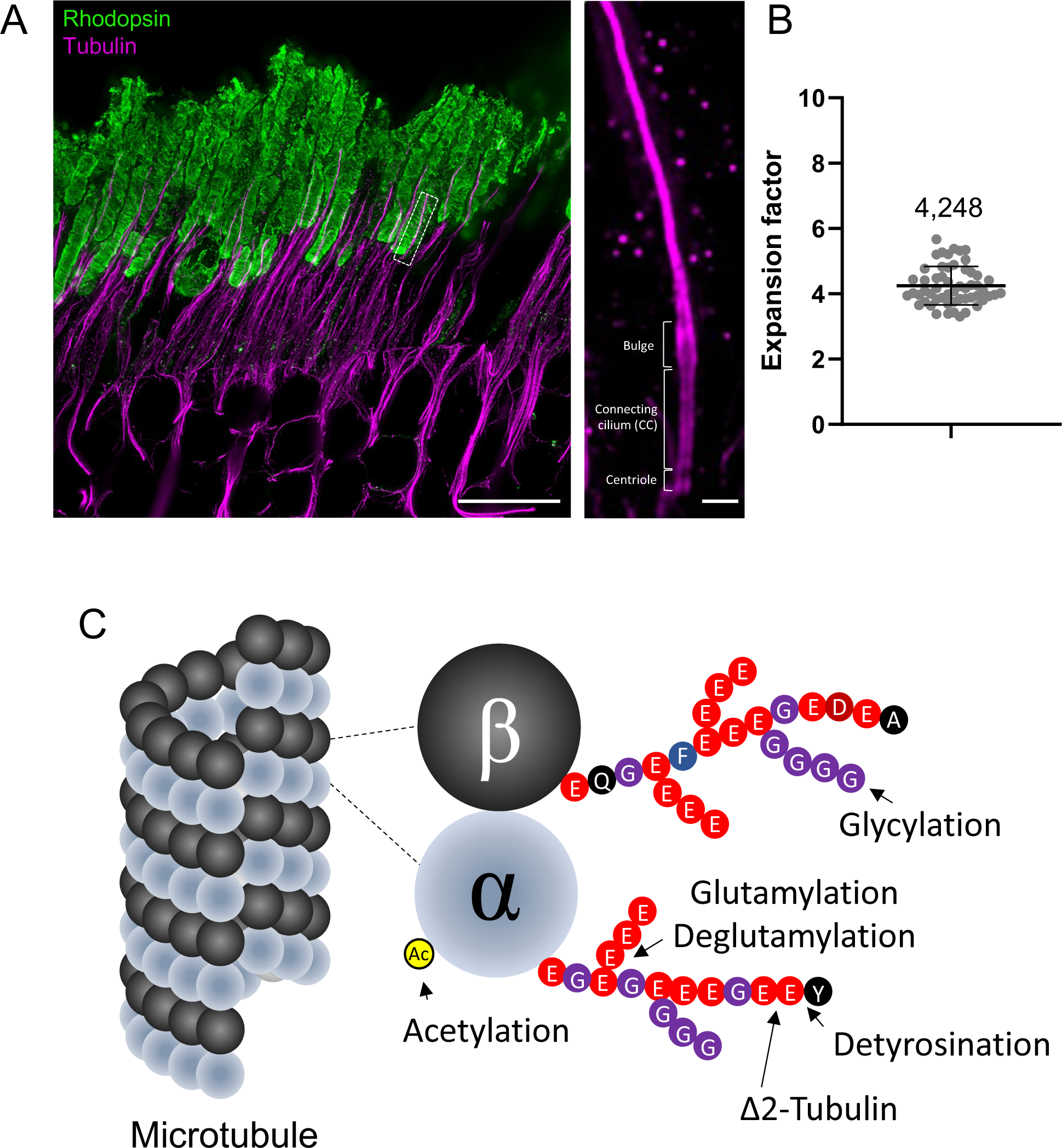
Architecture of the photoreceptor Outer Segment. (**A**) Expanded photoreceptor cell layer of a WT mouse retina stained for tubulin (magenta) and rhodopsin (green). On the right, inset of a single photoreceptor cell outer segment stained for tubulin, revealing the different regions of the cilium. Scalebars: left: 10µm; right: 500nm (**B**) Quantification of the expansion factor (EF) used for the whole study. EF = 4.248 +/- 0.588 (Mean +/- SD) (N>5) (**C**) Model explaining the tubulin code and highlighting the PTMs analyzed in this study.

**Supplementary Figure 2:**
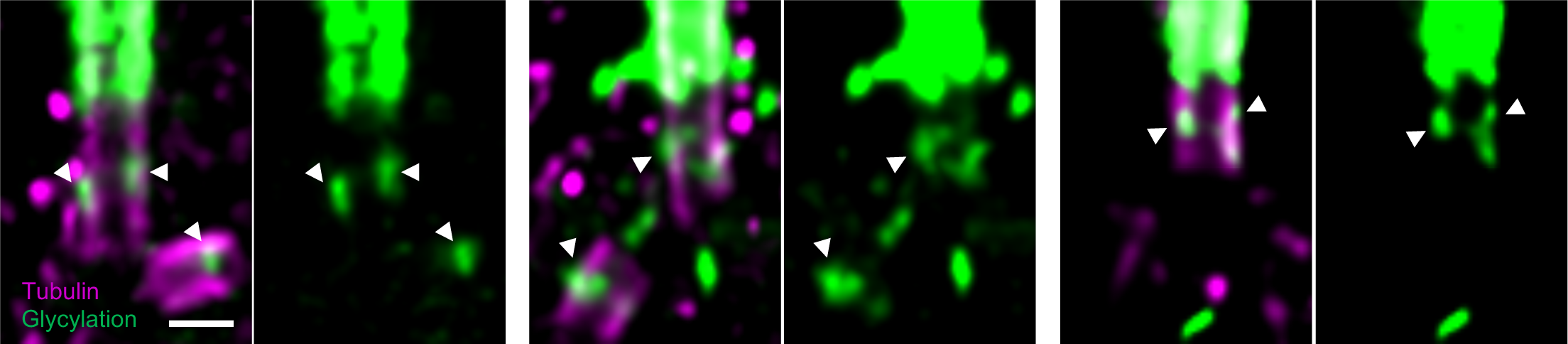
Glycylation signal at the level of the photoreceptor basal body. Expanded photoreceptor cell centrioles stained for TAP952 (green, saturated) and tubulin (magenta). Glycylation is observed at the level of centriole subdistal appendages (white arrowheads) when the TAP952 signal is pushed. Scalebar: 200 nm

**Supplementary Figure 3:**
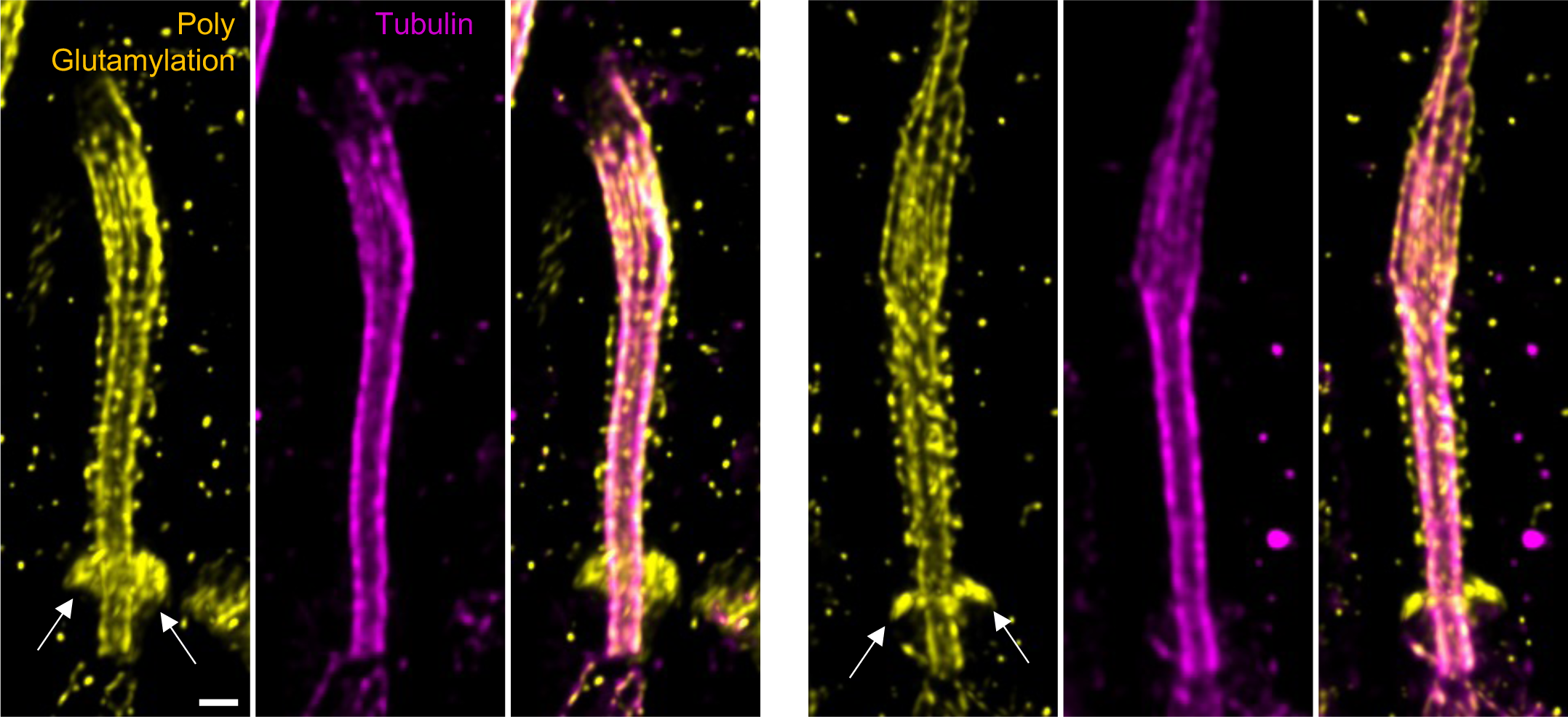
Polyglutamylation signal enrichment at the ciliary base. Expanded photoreceptor cell outer segments stained with PolyE (yellow) and tubulin (magenta). Polyglutamylation is sometimes observed enriched at the base of the cilium, similar to IFT trains formation. Scalebar: 200 nm

**Supplementary Figure 4:**
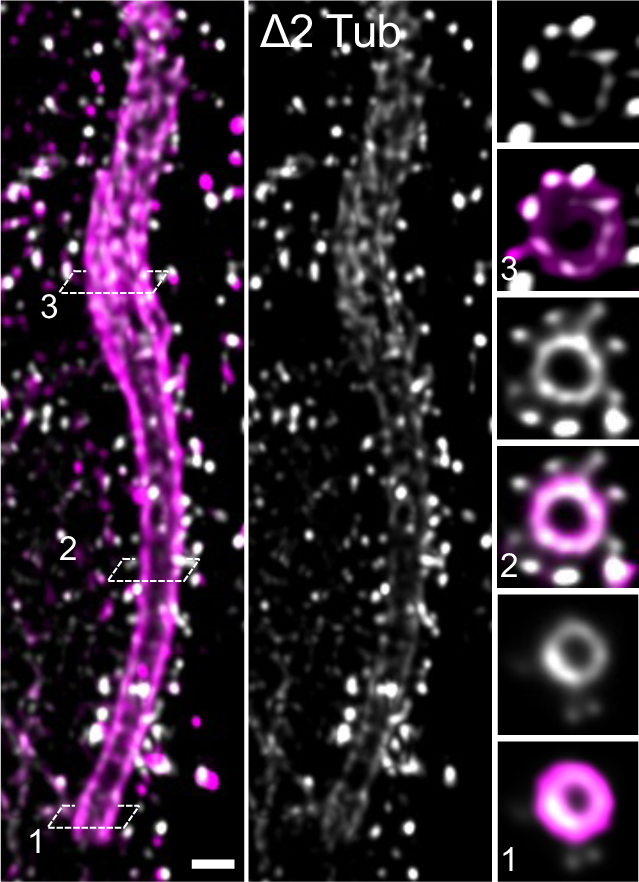
Delta2 tubulin signal in the OS. Molecular mapping of the delta2 tubulin (gray) along the photoreceptor outer segment axoneme stained for tubulin (magenta). Transversal section images corresponding to different regions of the OS (centriole (1), connecting cilium (2) and bulge (3), depicted by the dashed lines and numbers on longitudinal images) are represented on the right side. Scalebar: 200 nm.

**Supplementary Figure 5:**
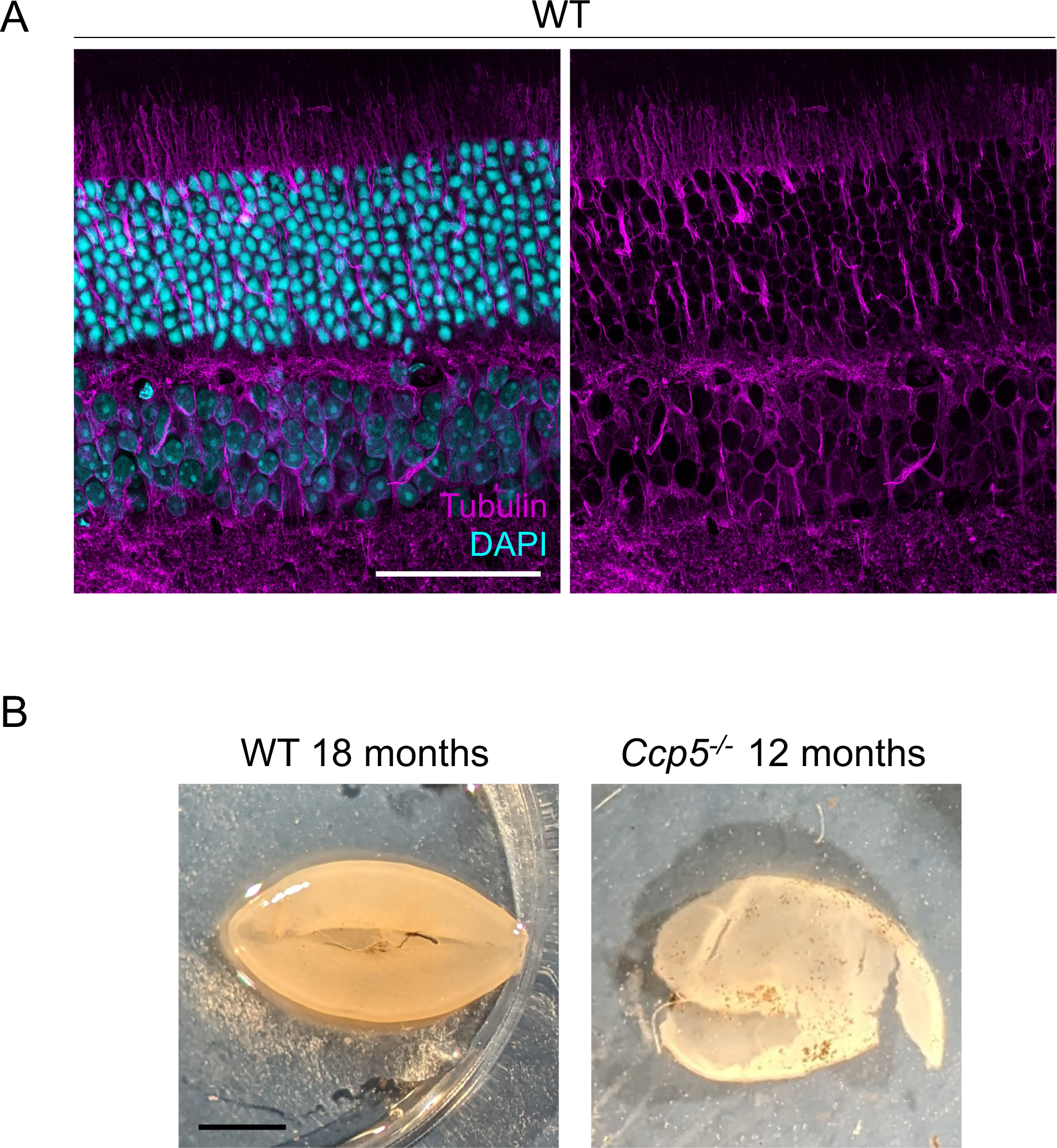
Global morphology of WT and *Ccp5^-/-^* retinas. (**A**) Expanded WT mouse retina stained for tubulin (magenta) and DAPI (cyan). Note that ONL thickness can be measured only with tubulin staining, where nuclei position is clearly visible. Scalebar: 50 µm. (**B**) 12 month-old WT (left) and *Ccp5^-/-^* (right) retinas during dissection. Note that *Ccp5^-/-^* retina is thinner and pigmented, reflecting a strong degeneration, a feature that we already observed previously (Faber et al., 2023). Scalebar: 1mm.

**Supplementary Figure 6:**
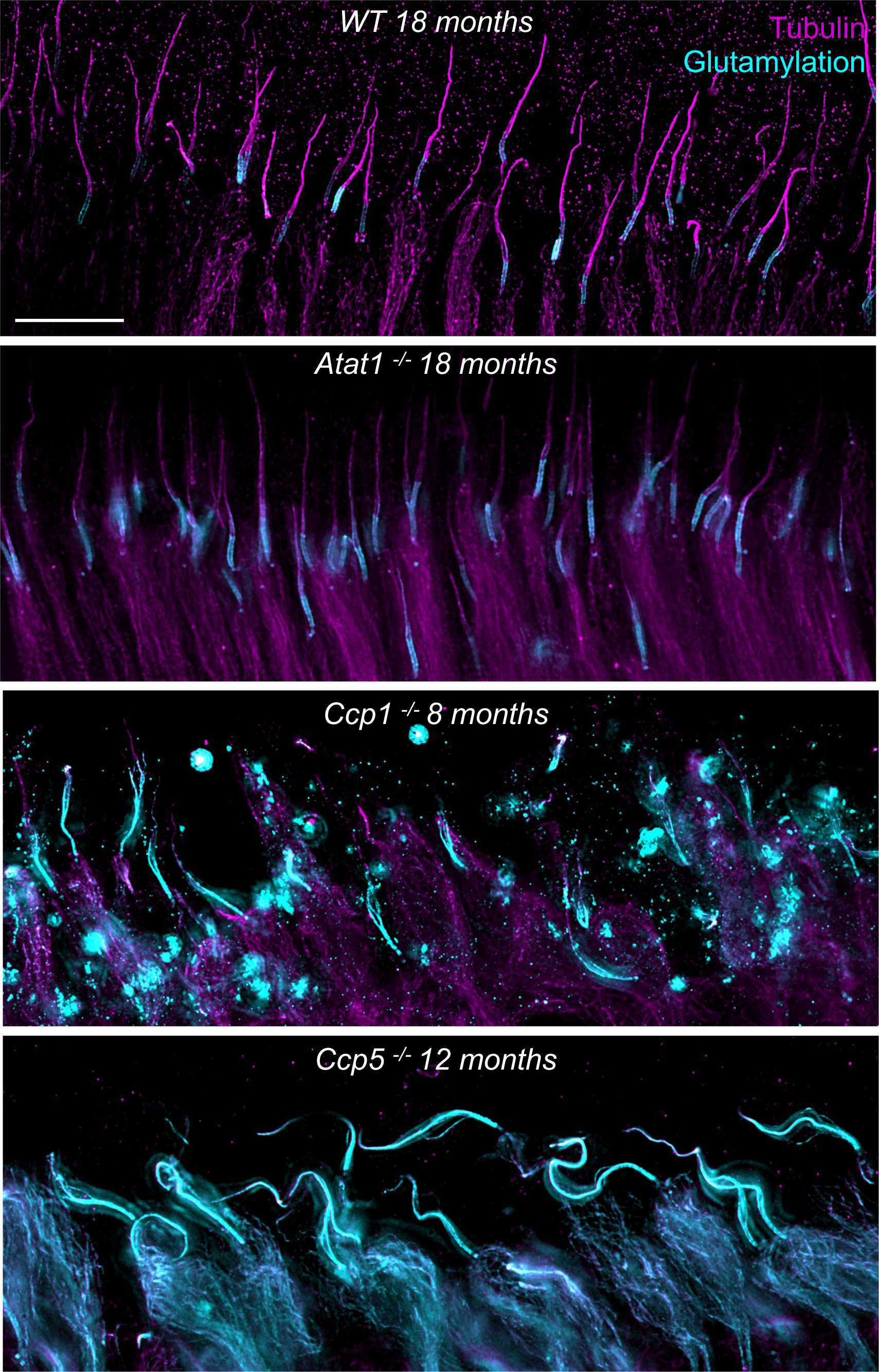
Glutamylation level observed in several PTM mutants. Large field of view of WT, *Atat1^-/-^*, *Ccp1^-/-^* and *Ccp5^-/-^* expanded photoreceptor cell outer segments stained with GT335 (cyan) and tubulin (magenta). Note the intense glutamylation signal inside photoreceptor cell bodies in *Ccp5^-/-^*retina. Scalebar: 5 µm.

**Supplementary Figure 7:**
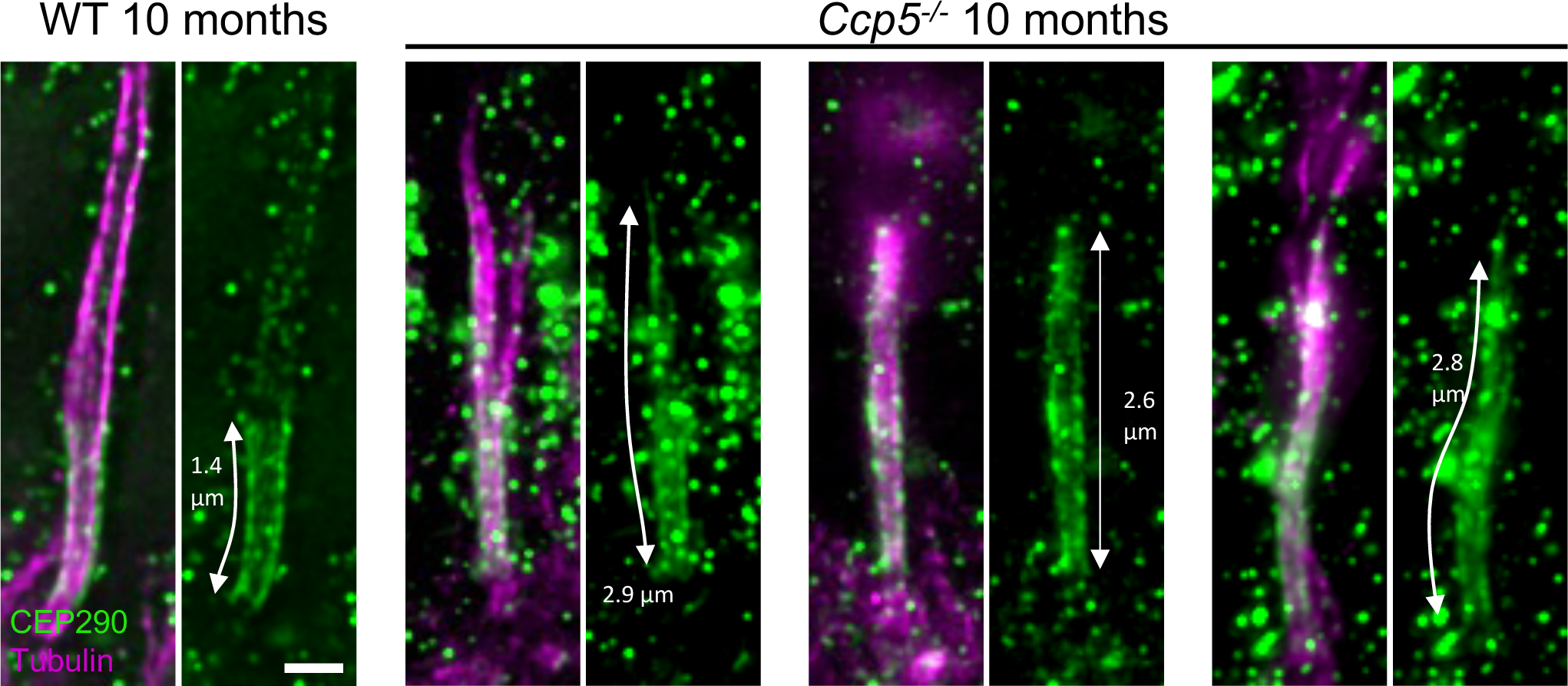
Exacerbated CEP290 signal in *Ccp5^-/-^* photoreceptor cells. Examples of 10-month-old WT or *Ccp5^-/-^* degenerating photoreceptor cells stained with CEP290 (green) and Tubulin (magenta) to highlight that inner and outer components of the CC are both exacerbated in the mutant retina. White double arrows reveal the length of CEP290 signal with the measurement corrected by the EF. Scalebar: 500 nm.

**Supplementary Figure 8:**
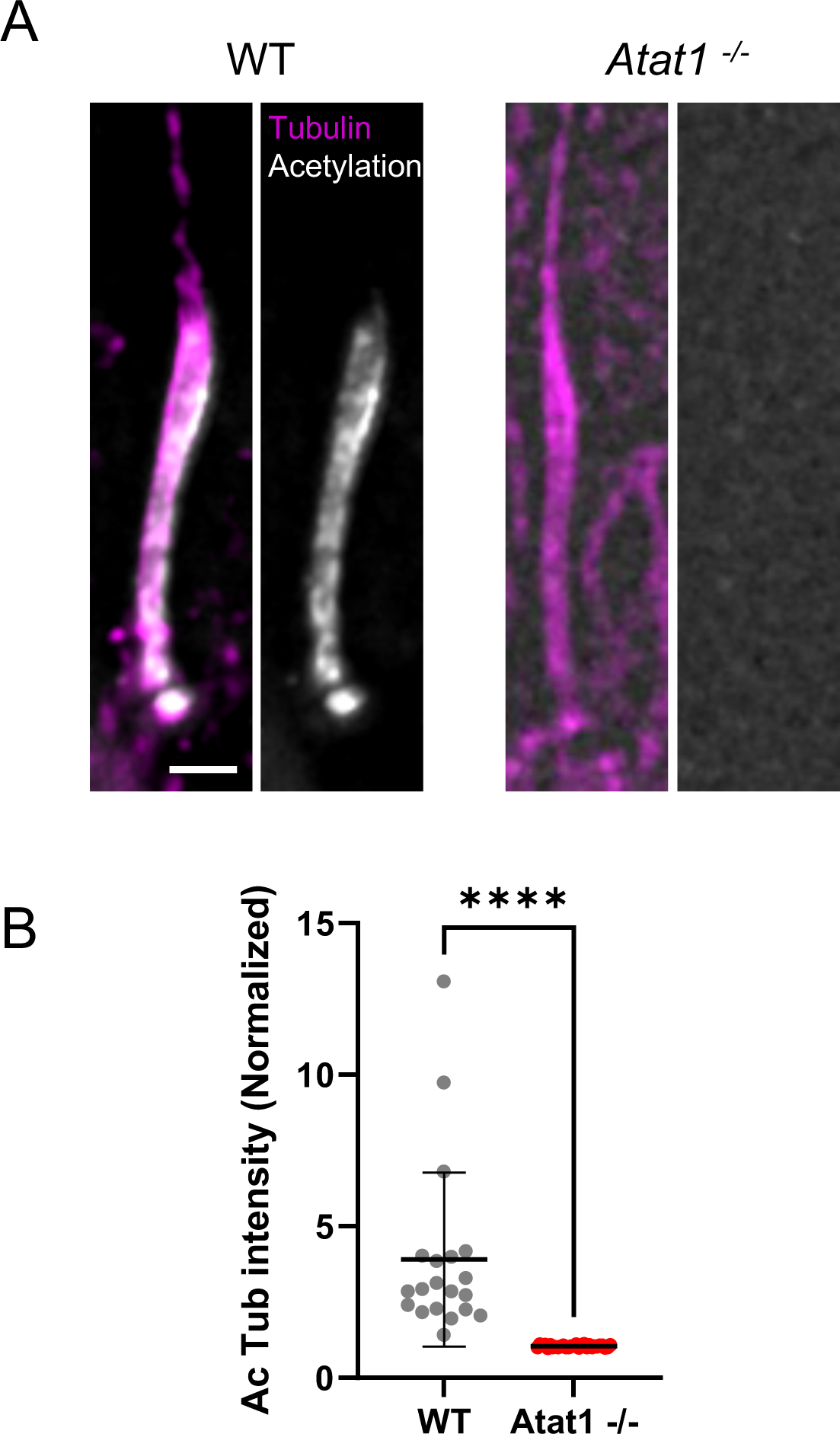
Loss of acetylation in the OS in *Atat1^-/-^* mice. (**A**) Expanded WT or *Atat1^-/-^* photoreceptor cells demonstrating the total loss of acetylated tubulin staining (gray), together with tubulin (magenta). Scalebar: 500 nm. (**B**) Quantification of the acetylated tubulin signal in WT or *Atat1^-/-^* photoreceptor cells. WT: 3.9 +/- 2.9; *Atat1^-/-^*: 1.04 +/- 0.04 (Mean +/- SD). N=3.

**Supplementary Figure 9:**
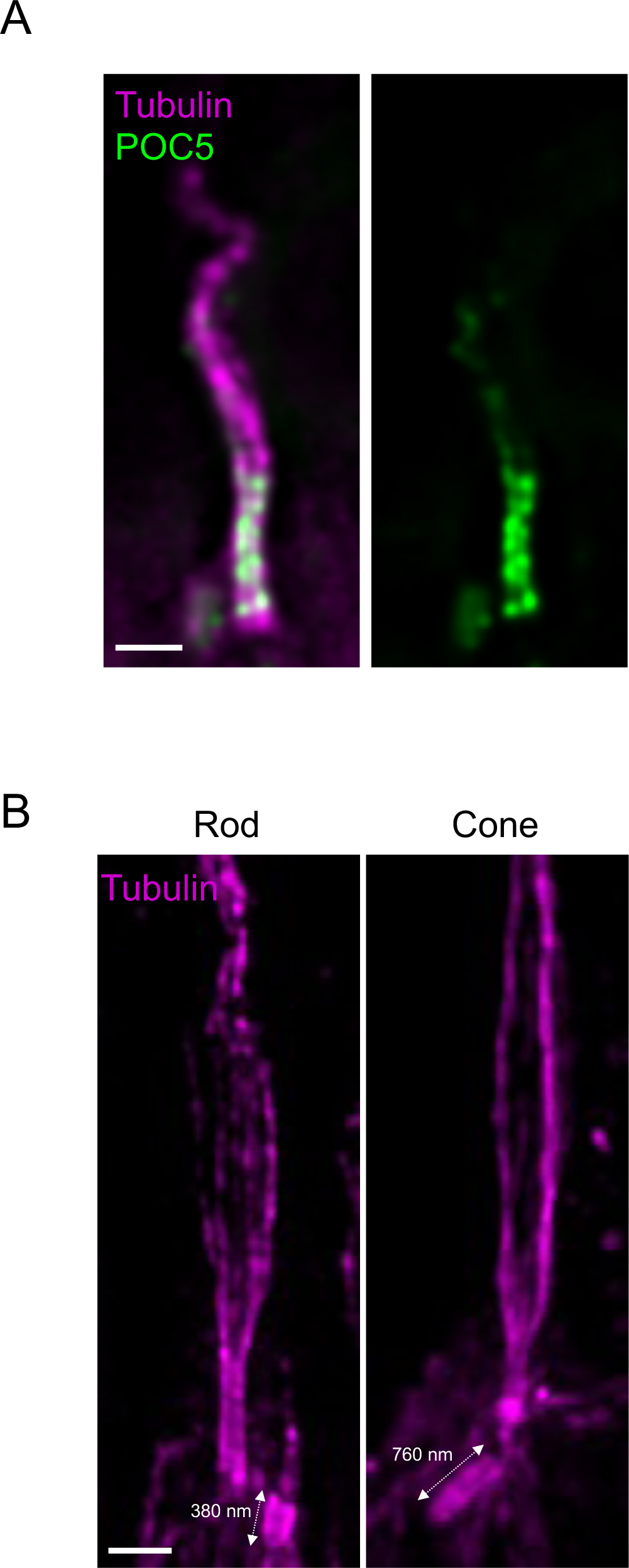
Connecting cilium inner scaffold in human photoreceptor cells. (**A**) Expanded human rod stained for POC5 (green) and tubulin (magenta). (**B**) Expanded human rod (left) or cone (right) photoreceptor cells stained for tubulin (magenta). Size of the daughter centrioles are highlighted by the white double headed arrows. Scalebar: 500 nm.

**Supplementary Figure 10:**
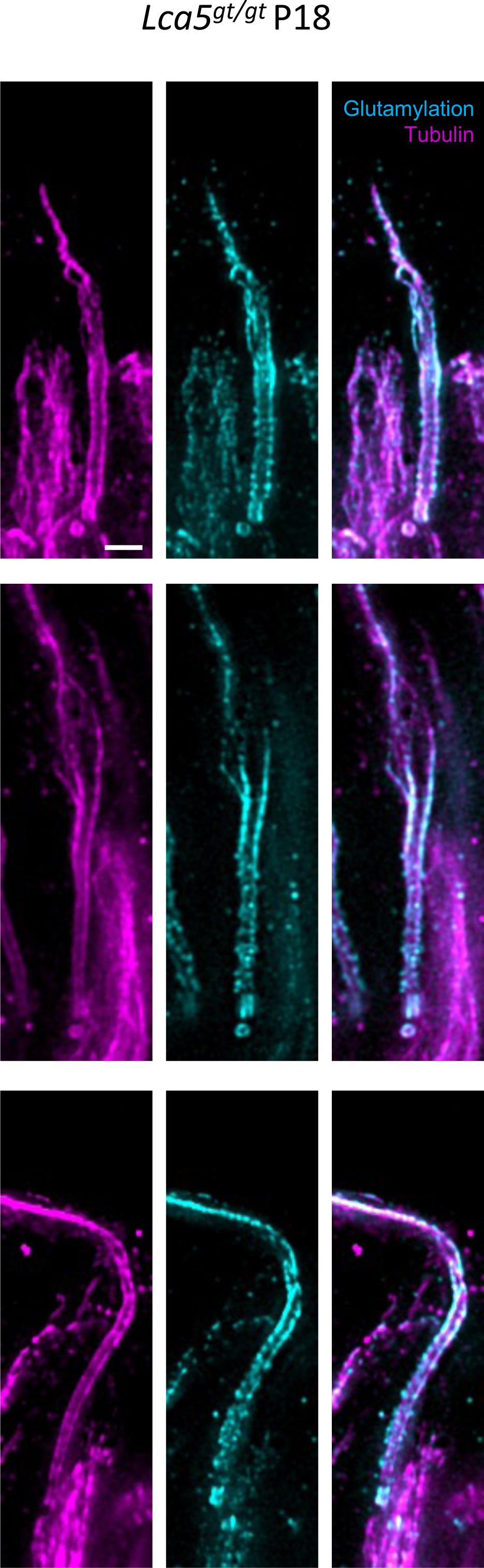
Hyperglutamylation of the OS in *Lca5^gt/gt^* photoreceptor cells. Expanded P18 *Lca5*^-/-^ photoreceptor cells stained for glutamylation (GT335, cyan) and tubulin (magenta). Scalebar: 500 nm

